# Lifespan Trajectories of the Brain’s Functional Complexity Characterized by Multiscale Sample Entropy

**DOI:** 10.1101/2025.03.03.641327

**Authors:** Dilmini Wijesinghe, Kirsten Lynch, Leon Askman, Dean C. Delis, Danny JJ Wang, Kay Jann

## Abstract

Resting state functional magnetic resonance imaging (rs-fMRI) is a widely used imaging modality that can capture spontaneous neural activity of the brain. The human brain is a complex system, and emerging evidence suggests that the complexity of neural activity may serve as an index of the brain’s capacity of information processing. In this study we used multiscale sample entropy (MSE) to analyze the complexity of rs-fMRI timeseries of 504 healthy subjects between the age of 6 to 85 years. We constructed the global and regional trajectories of brain’s functional complexity over the lifespan and analyzed its correlation with executive function. We observed a nonlinear trajectory of fMRI-complexity over the lifespan with a peak age occurring at 23 (95% CI 21.27, 26.38 years) years of age. Males reach the peak age of complexity at 26 years (95% CI 19.95, 33.14 years) whereas females at 23 (95% CI 20.16, 29.18 years). We found significant correlations between complexity and Number-Letter Switching of Trail Making Test in parietal and medial temporal lobes while Inhibition and Inhibition/Switching of Color Word Interference Test also revealed a significant negative correlation with rs-fMRI complexity in lateral and medial frontal cortex. These results help to understand behavior of fMRI-complexity with aging and reveals association with executive function of the brain. As a non-invasive biomarker, fMRI-complexity could provide novel approach to understand information processing capacity in the brain and deficits thereof in illness.

## 1. Introduction

Brain maturation is a complex and protracted process of progressive and regressive events that plays a critical role in cognitive development. During this process an individual undergoes a variety of significant changes physiologically, psychologically, and biologically. Understanding normal developmental and aging processes across the lifespan becomes critically important to lay the foundation for the detection, prevention and treatment of many brain disorders related to aging, and neurodevelopment disorders that may have lifelong impact ^[1,2]^. Establishing typical developmental trajectories is essential for detecting deviations indicative of psychiatric and neurological disorders. Such deviations may be crucial in predicting age-related cognitive decline and in identifying biomarkers that enable the early detection of neurodevelopmental disorders and neurodegeneration ^[3]^.

The human brain consists of neurons, synaptic connections and networks of neuronal populations that have hierarchical relationships and give rise to nonlinear, dynamic, and complex behaviors. Over the years, efforts in understanding the development and aging of the human brain have employed an array of different modalities. Most prevalent are structural magnetic resonance imaging (MRI) methods to assess metrics such as cortical thickness and volume or diffusion weighted imaging to assess maturation of white matter fiber tracts ^[4,5]^. Resting-state fMRI (rs-fMRI) has been used to investigate inherently fluctuating signals and their interactions between different brain regions ^[6]^. Functional connectivity analysis of fMRI data revealed that the brain uses stable networks to process the information it receives from the external environment ^[7]^.

In information theory, information processing capacity (uncertainty of information) can be measured using statistics-based entropy measures. These methods are nonlinear and dynamic measures and can be applied to analyze complexity in time series data. These techniques have been utilized in analyzing resting-state fMRI timeseries data and obtaining quantifiable outcomes for complexity of the brain signal recordings and are complementary to common functional connectivity approaches. One popular interpretation of this measure of brain complexity is that it measures fluctuations among brain networks since entropy is a measure of number of ways a system can be configured ^[8]^. Novel complexity metrics have been introduced over the recent years and they were incorporated into evaluating fMRI signal complexity as a metric reflecting regional information processing capacity. Therefore, we can hypothesize that brain complexity measures the brain’s ability to change, process and adapt (flexibility) information received by switching between networks and network configurations.

Over the past decade there are fMRI studies that analyzed complexity with age using different complexity metrics such as fractal dimensions or entropy methods ^[9,10,11]^. Overall, these studies observed a decline in complexity with age. Besides lifespan trajectories of normal maturation, there is evidence from divergent neurodevelopment and aging supporting the notion that brain entropy can detect disease related deviations from normal trajectories ^[12,13,14,15]^. Also, according to the hypothesis, brain entropy can be seen as a measure of transitions and communication within and between networks. Cognitive abilities such as adaptive responses, reasoning, learning, problem solving and planning which are associated with intelligence would require accessing large number of brain states to make appropriate integrations ^[8]^. This raises the question whether complexity emerging from neuroimaging is reflecting general intelligence or cognitive ability in certain domains. To answer these questions, additional studies are necessary to explore the underlying physiological meaning of complexity ^[16]^, its normal maturational trajectories, and its relation to cognitive abilities.

In this study we investigate the changes in Multiscale Sample Entropy ^[17]^ (MSE) over the lifespan along with sex effects. MSE is an improved metric of Sample Entropy (SampEn) which calculates the rate of generation of new information in dynamical process ^[18,19]^. MSE offers an appealing complexity metric for fMRI time series data analysis, as it can differentiate random noise from complex processes with self-similarity ^[18,20]^. Additionally, the temporal complexity of rs-fMRI measured by MSE was demonstrated as reproducible ^[19]^. However, MSE has yet to be used to study development and healthy aging over the entire lifespan using rs-fMRI. Furthermore, we investigate the association between lifespan trajectories of fMRI-complexity and age dependent changes in executive cognitive function. Executive cognitive function refers to higher order cognitive abilities and is generally associated with frontal lobe function ^[21]^.

## 2 Materials and Methods

### 2.1 Study Characteristics and MRI Acquisition

The study included data from the Nathan Klein Institute (NKI) – Rockland Sample. Anatomical as well as rs-fMRI data of 526 subjects with age ranging from 6 to 85 years of age were included. FMRI data were acquired using a gradient-echo (GRE) EPI sequence with the following parameters: TR/TE = 1400ms/30ms; 2 mm isotropic resolution; 64 slices, 112×122 matrix size, FA = 65°, total acquisition duration = 10mins. Also, for T1w images, TR/TE = 1900ms/2.52ms; 1mm isotropic resolution; 176 slices, 256×256 matrix size, FA = 9°, total acquisition duration 4 minutes and 18 seconds. The enhanced NKI project was aimed at creating a large scale (number of participants > 1000) community sample of participants across the lifespan ^[22,23]^.

### 2.2 Data Pre-processing

Data pre-processing and denoising was performed in CONN-toolbox and included image realignment, regression of motion parameters and derivatives, regression of cerebrospinal fluid and white matter (WM) signal fluctuations using a-CompCorr, linear drift removal using a high-pass filter, coregistration to individual T1w images and normalization to MNI template space. Twenty-two subjects with composite motion score greater than 0.5 mm were excluded from the study. Composite motion score was obtained from the ‘art_regression_outliers_and_movement’ file from the CONN toolbox. The remaining total of 504 subjects were divided into 5 age groups using k-means clustering (Table 1): Age 6-13 (Pre-Puberty, 19 Females, 22 Males), 14-25 (Adolescence, 68 Females, 64 Males), 26-38 (Young Adult), 39-63 (Middle Age, 32 Female, 43 Males), 64-85 (Senior, 67 Females, 35 Males). Resting state fMRI-complexity was estimated by means of MSE using an in-house complexity toolbox, LOFT Complexity Toolbox (github.com/kayjann/complexity).

**Table 1:** Age groups.

| Group | Age (years) | Number of Female/Male Participants | Total Number of Participants |
| --- | --- | --- | --- |
| Pre-Puberty | 6 - 13 | 17/17 | 34 |
| Adolescence | 14 - 25 | 66/64 | 130 |
| Young Adult | 26 - 38 | 32/28 | 60 |
| Middle Age | 39 - 63 | 141/39 | 180 |
| Senior | 64 - 85 | 65/35 | 100 |

### 2.3 Sample Entropy (SE)

Sample entropy ^[18]^ is a variant of Approximate Entropy (ApEn) ^[24]^ that was developed for short and noisy physiological signals. Sample entropy for a given two embedding dimensions can be calculated using

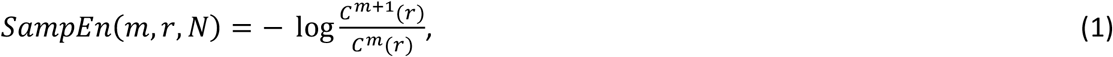

where *m* (pattern matching length) denotes the length of block or pattern to be compared, *r* (pattern matching threshold) is the distance threshold and *N* is the length of the timeseries. The parameters *m* and *r* must be chosen and *C*^*m*^(*r*) term in Eq (1) calculates the average conditional probability of recurring data points within the chosen pattern matching threshold ^[20]^.

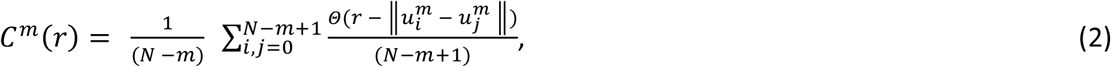

where 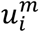 and 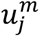 are two embedding dimensions with pattern matching length m and Θ is the Heaviside function.

### 2.4 Multiscale Sample Entropy (MSE)

Sample entropy evaluates complexity of a timeseries on a single scale whereas MSE evaluates complexity by using multiple coarse-grained timeseries ^[17]^. This aids to filter out the high frequency fluctuations ^[20]^. A new coarse grained timeseries is created by averaging consecutive time points of the original timeseries within non-overlapping windows. For a given timeseries {*x*_1_, …, *x*_*N*_}, averaged time point 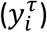 of the new timeseries given by

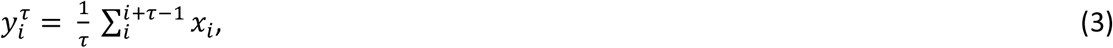

where τ is the time scale which determines the number of consecutive time points and length of the new timeseries is *N*/τ. Accordingly scale 1 is the original timeseries and its frequency (Hz) is defined as 1/TR. Subsequent higher scales will thus have lower frequencies defined as 1/(scale*TR).

MSE was calculated as the area-under-the-curve across SampEn up to τ = 13 scales, with pattern matching length m = 2 and pattern matching threshold r = 0.3 ^[25,26]^. Complexity maps were segmented into gray matter (GM) and white matter (WM) tissue compartments, and the maps were further parcellated into 133 cortical gray matter regions of interests (ROIs) based on the Harvard Oxford atlas. Brain stem, cerebellum and cerebellar vermis regions were excluded, and complexity of 105 cortical brain regions were analyzed. Average MSE of GM voxels and the 105 cortical regions of interest (ROIs) were calculated for each subject.

### 2.5 Statistical Analysis

We evaluated trajectories for global GM complexity as well as regional trajectories of rMRI-complexity and further investigated sex differences in these trajectories. To determine the trajectory of the global GM complexity changes across the lifespan, we applied five models for Bayesian Information Criterion (BIC) based model selection: linear, quadratic, cubic, quartic, and double exponential functions. The BIC was calculated using MATLAB to determine the best model fitting for the data. Peak age of complexity maturation in whole, female, and male groups were determined by running stratified bootstrapping analyses for 1000 iterations with 20 age bins for stratification. The BIC model selection was applied in every iteration to determine the best fit model. Bootstrap iterations that do not successfully model the data using aforementioned list of polynomials and data with linear distribution from age 6 were ignored during calculations. Confidence intervals (percentile method) and mean of peak age of complexity maturation were determined using the peak age distributions obtained from bootstrapping. A similar approach was applied to regional analyses to assess the best fit model for the data distribution of each ROI by applying the five models: linear, quadratic, cubic, quartic, and double exponential functions along with BIC. Peak age of complexity maturation was calculated using the fitted models. Rates of increase and decrease were calculated by dividing datasets into two: pre-peak data and post-peak data. Using linear models, slope percentages 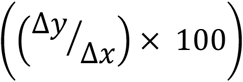 before and after the peak were calculated. Finally, K-means clustering was applied to cluster the peak age of complexity maturation and slopes of increase and decrease of complexity over the lifespan to reveal patterns or networks with similar trajectories. All analyses were repeated by separating the cohort into female and male to assess sex-specific trajectories. Three-dimensional (3D) rendered images were created for data visualization of cortical areas using the Quantitative Imaging Toolkit (QIT) software ^[27]^.

To evaluate the potential impact of gray matter (GM) volume on this analysis we calculated the GM volume of each subject using FMRIB Software Library (FSL) and added GM volume as a covariate in a linear model of complexity vs age (see supplemental material). Statistical significance level is considered as *p* < 0.05 throughout the analyses with correction for multiple comparisons at alpha = 0.0083.

### 2.6 Delis-Kaplan Executive Function Scores (D-KEFS)

The NKI study participants underwent neurocognitive testing using Delis-Kaplan Executive Function System test (D-KEFS) ^[28]^. The D-KEFS is a neurophysiological test which is widely used on testing cognition and executive function and consists of 9 stand-alone tests: Color-word Interference Test (CWIT), Design Fluency Test (DFT), Proverb Test (PT), Sorting Test (ST), Tower Test (TT), Trail Making Test (TMT), Verbal Fluency Test (VFT), Word Context Test (WCT), and Twenty Questions Test (TQT). After careful consideration of clinical applications of D-KEFS we have chosen 6 primary tests for the analysis: TMT (Number-Letter Switching), VFT (Letter Fluency and Category Switching Total: Number of Correct Switches), CWIT (Inhibition and Inhibition/Switching) and DFT (Switching). TMT aims at assessing set-shifting, attention, and processing speed and visuomotor scanning ^[29,30]^. VFT aims at assessing category fluency, set-shifting and letter fluency ^[29]^. CWIT measures are effective on accessing working memory, inhibitory control, cognitive flexibility, semantic activation, and response selection ^[29]^. DFT aims at assessing planning/initiation, cognitive flexibility/divergent thinking, and fluency in generating visual patterns ^[29,31]^.

After comparing subject IDs with phenotypic data and neuroimaging data, 494 participants had sufficiently complete cognitive test score data. TMT-Number-Letter Switching and DFT-Switching had no missing values while in the remaining four tests altogether had 8 missing values and those missing values were imputed using spline interpolations. The continuous values in the six tests were standardized to z scores. Statistical analysis between complexity and cognitive test scores in global and regional data were calculated by Spearman’s rank correlation coefficient. Furthermore, a one-way ANOVA followed by post-hoc tests were applied on D-KEFS scores separated by age groups in Table 1.

## 3 Results

### 3.1 Trajectories for whole brain gray matter functional complexity

For the global trajectory of complexity, the lowest BIC was observed for the double exponential function. The peak of the global trajectory of mean MSE occurs at the age 23.76 years ([21.27, 26.38 years], CI 95%). Before the age of 23, complexity and age have a position correlation coefficient r = 0.3418 (p = 5.653 s 10^−5^). After the age 23, the correlation coefficient was r = −0.247 (p = 2.170 × 10^−6^).

Regarding sex effect, the complexity peak in females occurs at the age of 23.67 ([20.16, 29.18 years], 95% CI) years whereas for males, the peak age occurs at 26.07 years ([19.95, 33.14 years], CI 95%) (Figure 1). Generally, after the age of 23 years in the young adult, middle aged and senior groups there is a steady decline in complexity with age. There was no significance effect of GM volume on mean MSE, and the recorded p value was 0.995 (see supplemental material).

**Figure 1.**
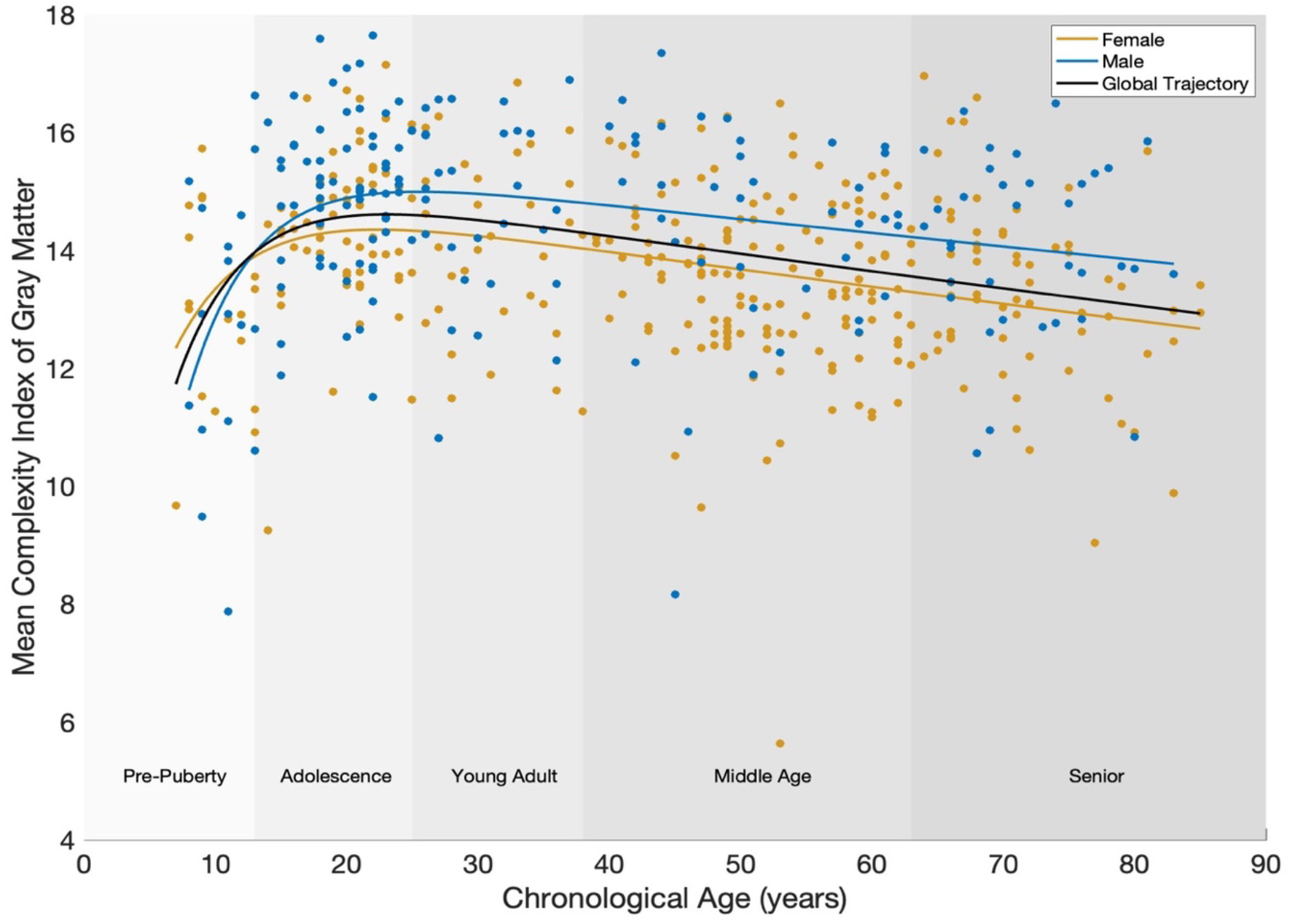
Global trajectory of complexity (Black). Fitted double exponential function for complexity trajectory of female and male given by yellow and blue lines. Mean complexity index of gray matter less than 7 were excluded from the polynomial fitting.

### 3.2 Peak age of complexity

Qualitative observation indicates that frontal and prefrontal cortical areas have a later peak age than parietal cortical areas. In the whole sample, the peak age of complexity in most areas in frontal and parietal lobe reflecting higher association cortex occurs after 22 years (Figure 2a), while occipital and temporal areas showed peak age before 20 years. This pattern was similar and even more pronounced in the female group (Figure 2b) while the male group showed overall later peak ages (Figure 2c). Notably, the peak age of complexity among female participants varies from 19.4 to 33.7 years, whereas for male participants, it varies from 21.1 to 34 years.

**Figure 2.**
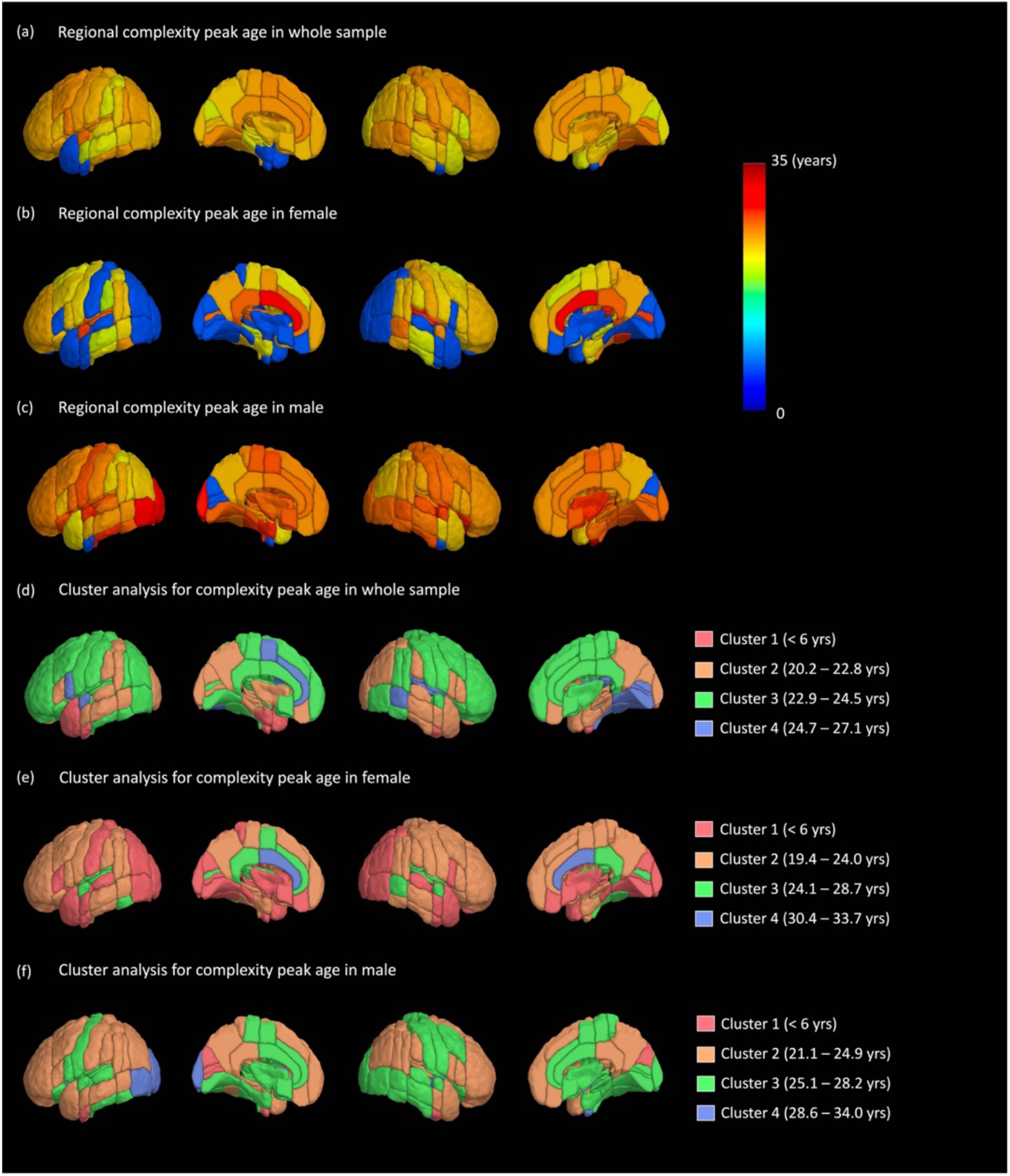
Regional peak age when maximal cortical complexity is measured in global (a), women (b) and men (c). Clusters of peak age of complexity maturation in global (d), women (e) and men (f). Complexity peak age of women in cortical areas varies from 19.4 to 33.7 years (e). In men this range varies from 21.1 to 34 years (f). Cluster 1 represents peak age for ROIs that have a linear decline from age 6.

To identify distinct cortical areas that undergo similar development and decline, we used clustering analysis to cluster cortical areas according to their fMRI-complexity profiles across the whole cohort, and separated groups by sex. We chose to set the cluster number to four similar to previous reports of distinct spatiotemporal development of cortical areas ^[32]^.

Figure 2d of whole sample cluster analysis shows left hemisphere temporal lobe with complexity peak age under-6 years. Late peak age of complexity is observed in most of the areas of frontal lobe and anterior cingulate gyrus.

In the female group, the complexity peak in regions of occipital pole and temporal pole falls into the under-6 years age group, as the observed data trend was linear from age 6 to 85 years (Figure 2e). In males, however left hemisphere occipital region reaches peak age of complexity later than other regions (28.6 – 34 years) (Figure 2f). Overall, both male and female group showed later peak of complexity in frontal cortex and anterior cingulate gyrus.

### 3.3 Rate of increase and decrease of complexity over the lifespan

Before the global peak at age 23, the rate of increase in complexity with age in frontal and parietal lobes are higher than the rest of the brain areas (Figure 3a). This observation prevails in male and female groups (Figure 3b & 3c).

**Figure 3.**
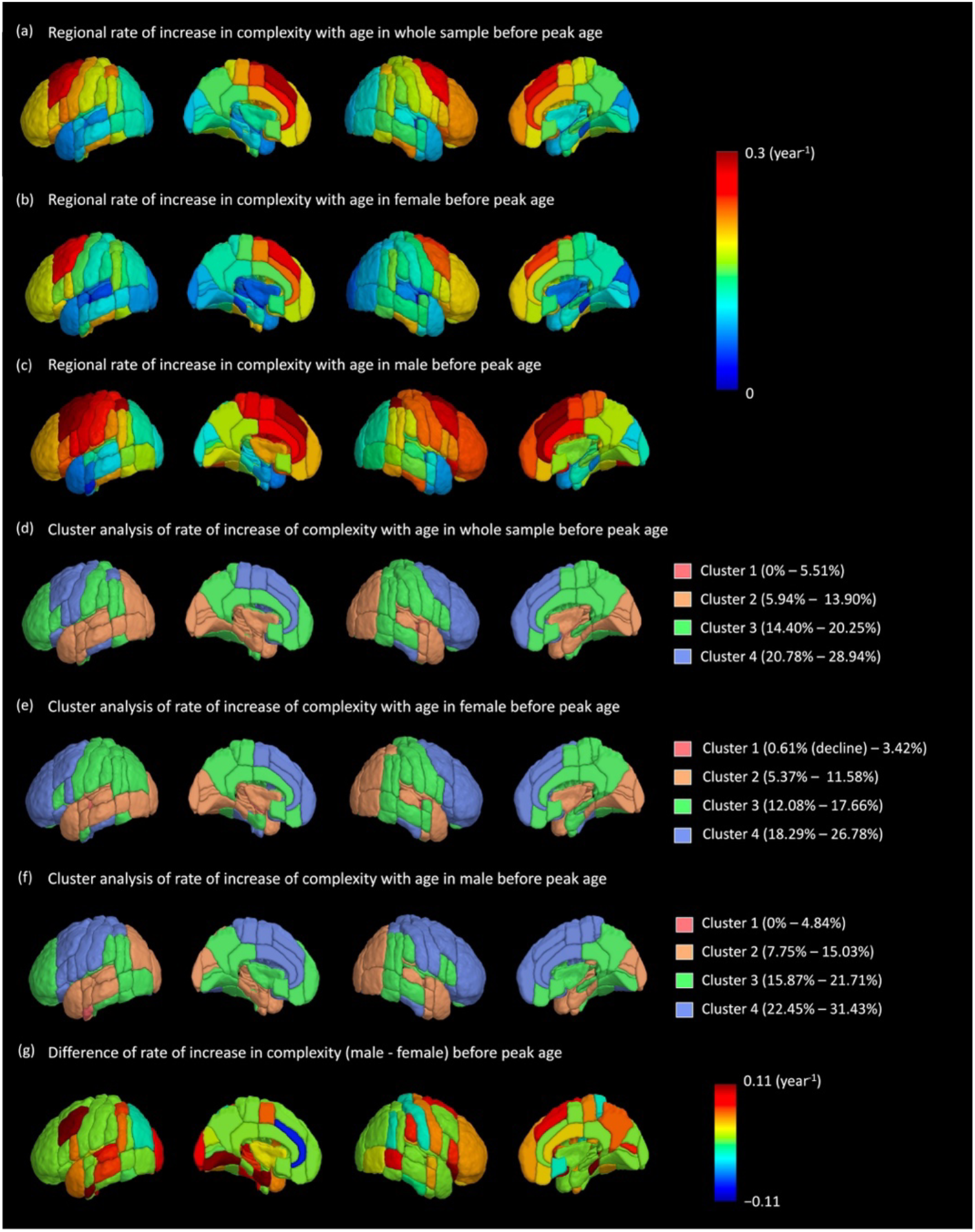
Change of cortical complexity before peak age. (a) Increase of cortical complexity with age (global). (b) Increase of cortical complexity with age in women. (c) Increase of cortical complexity with age in men. Clusters of rates of increase of complexity with age in global (d), women (e) and men (f). (g) Difference of rate of increase in complexity between male and female. Maximum and minimum values of slope percentage for each cluster are shown in the legend.

The cluster analysis clearly separates occipitotemporal (Cluster 2) from parietal (Cluster 3) and from frontal cortex (Cluster 4). These clusters define different rates of increase in adolescents to early adulthood with Cluster 4 showing more rapid increase than Cluster 3 and Cluster 2 respectively.

Frontal lobe of right hemisphere has the highest rate of increase in complexity compared to other regions in whole sample, female, and male groups (Figure 3d, 3e, 3f). In male group parietal lobe of left hemisphere has the highest rate of increase in complexity. In the female group, 104 ROIs showed positive slopes, except transverse temporal gyrus, which had a negative slope (Cluster 1, Figure 3e).

Overall, male group shows higher rate of increase in complexity compared to female group in all clusters (Figure 3c). Figure 3g illustrates the magnitude of the difference in slope between the male and female groups for each ROI (male - female). Except temporal pole and middle temporal gyrus, the male group has a higher rate of increase in complexity compared to female group (Figure 3g). However, these sex specific effects are not statistically significant except in superior parietal lobe (right), central opercular cortex (left) and thalamus (left).

After reaching a peak in complexity at age 23, moderate rate of decrease between 0 to 0.05 per year is shown in most of the brain regions in both whole sample and female group (Figure 4a and 4b).

**Figure 4.**
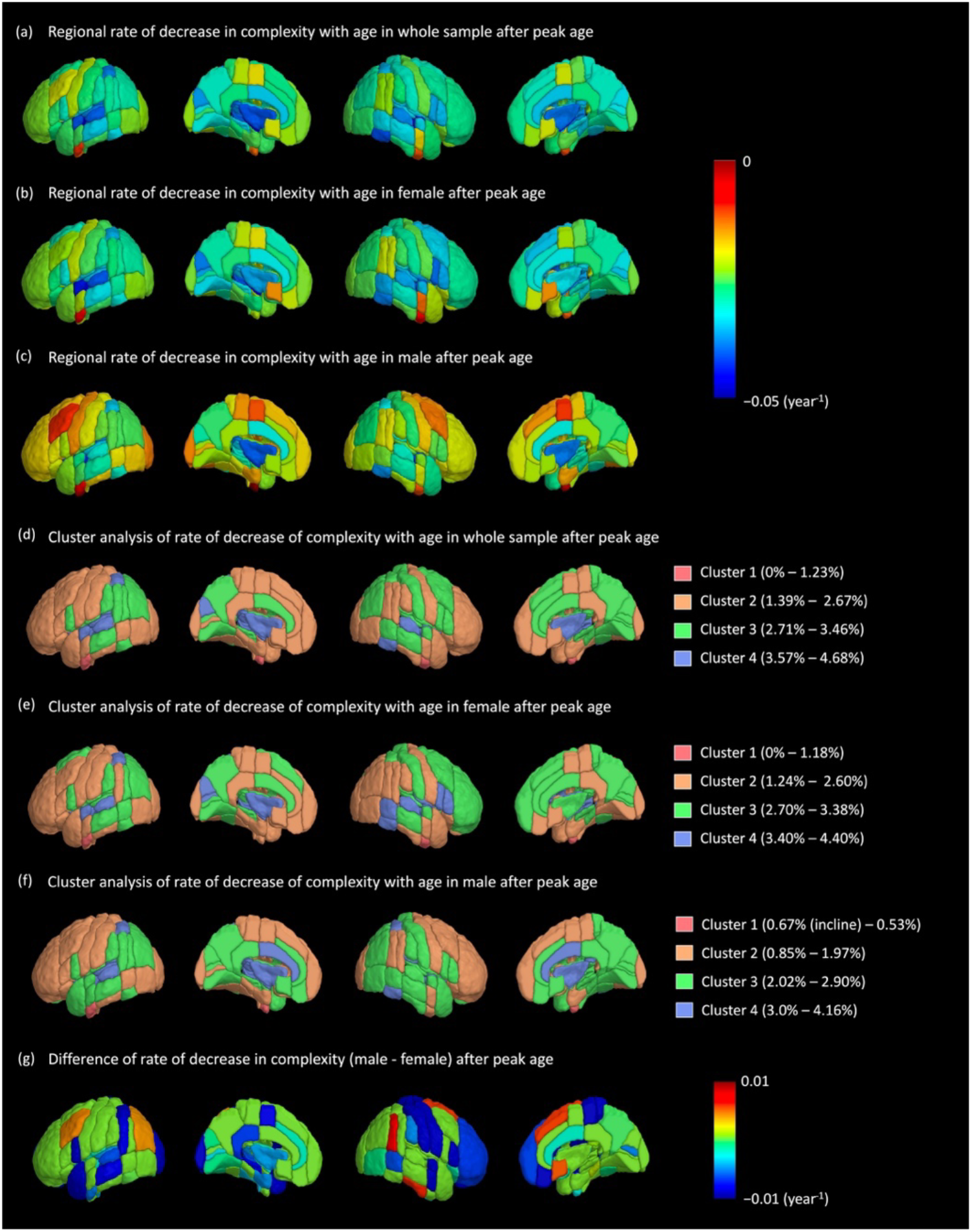
Change of cortical complexity after peak age. (a) Decrease of cortical complexity with age (global). (b) Decrease of cortical complexity with age in women. (c) Decrease of cortical complexity with age in men. Clusters of rates of decrease of complexity with age in global (d), women (e) and men (f). (g) Difference of rate of decrease in complexity between male and female. Maximum and minimum values of slope percentage for each cluster are shown in the legend.

However, in male group frontal and anterior parietal lobes show slower rate of decrease compared to rest of the regions (Figure 4c).

In cluster analysis prefrontal cortex and central gyri of left hemisphere showed rate of decrease of complexity between ~1% - 2.67% and this result is consistent in whole sample, female, and male groups (Figure 4d, 4e, & 4f). Female group showed faster rate of decline in frontal lobe of right hemisphere compared to left hemisphere male group (Figure 4e). In the male group, both right and left frontal lobe exhibit a slower decline compared to other brain areas (Figure 4f). Fastest rate of decline in female group is mainly in insular cortex and in male group anterior cingulate gyrus and insular cortex both have the highest rate of decline. Also, 104 ROIs in male group showed a negative slope, except for the temporal fusiform cortex, which had a positive slope (Cluster 1, Figure 4f). Overall, female group showed faster rate of decline in all clusters compared to male group. The magnitude of the difference in the rate of decrease between the male and female groups is illustrated in Figure 4g. Eighty-two percent of brain areas showed a higher rate of decline in complexity in females compared to males (Figure 4g).

In general, we observed a counterpose of results between male and female when comparing results before peak age and after peak age. These sex-specific observations on rate of decrease of complexity are not statistically significant.

### 3.4 Cognitive scores and complexity

We observed a strong significant negative correlation with global fMRI-complexity in TMT-Number-Letter Switching, VFT-Letter Fluency, CWIT-Inhibition and CWIT-Inhibition/Switching (Table 2). TMT-Number-Letter Switching, CWIT-Inhibition and CWIT-Inhibition/Switching has significant correlation with complexity. Other D-KEFS tests did not show significant correlation with complexity. The goal of the correlation analysis was to observe existence of a monotonic relationship between fMRI complexity and D-KEFS scores. Spearman’s rank correlation coefficient is suitable when data are heavy-tailed, include outliers ^[33]^, or to observe a monotonic relationship in continuous data. However, Pearson correlation of fMRI-complexity and D-KEFS were also calculated, and similar outcome of significance was observed (see supplemental material).

**Table 2:**
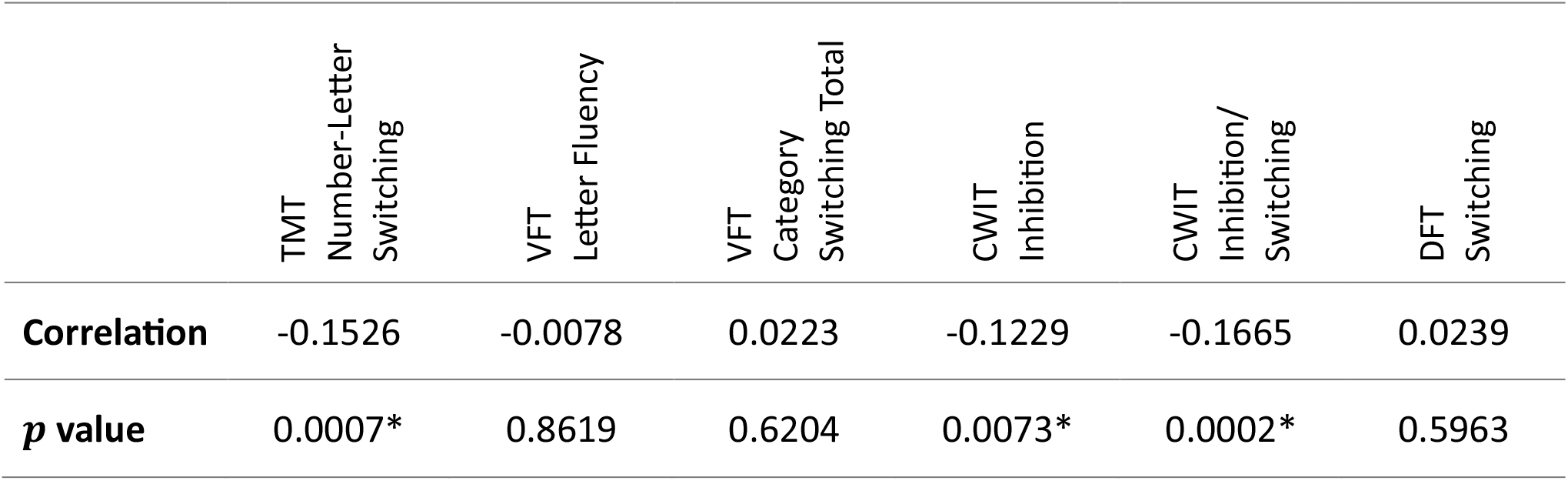
Spearman’s rank correlation between complexity and D-KEFS tests. (*significant after correction for multiple comparisons (adjusted *p* value 0.0083))

|  | TMT<br>Number-Letter<br>Switching | VFT<br>Letter Fluency | VFT<br>Category<br>Switching Total | CWIT<br>Inhibition | CWIT<br>Inhibition/<br>Switching | DFT<br>Switching |
| --- | --- | --- | --- | --- | --- | --- |
| **Correlation** | -0.1526 | -0.0078 | 0.0223 | -0.1229 | -0.1665 | 0.0239 |
| ***p* value** | 0.0007* | 0.8619 | 0.6204 | 0.0073* | 0.0002* | 0.5963 |

In regional analysis of fMRI-complexity and D-KEFS scores, TMT Number-Letter Switching showed significant negative correlation in parietal and medial temporal lobe (Figure 5a), whereas both CWIT Inhibition and CWIT Inhibition/Switching showed significant negative correlation with fMRI-complexity in lateral and medial frontal cortex (Figure 5b and5c). Among CWIT inhibition and CWIT inhibition/Switching, the later showed overall stronger negative correlation than CWIT inhibition.

**Figure 5.**
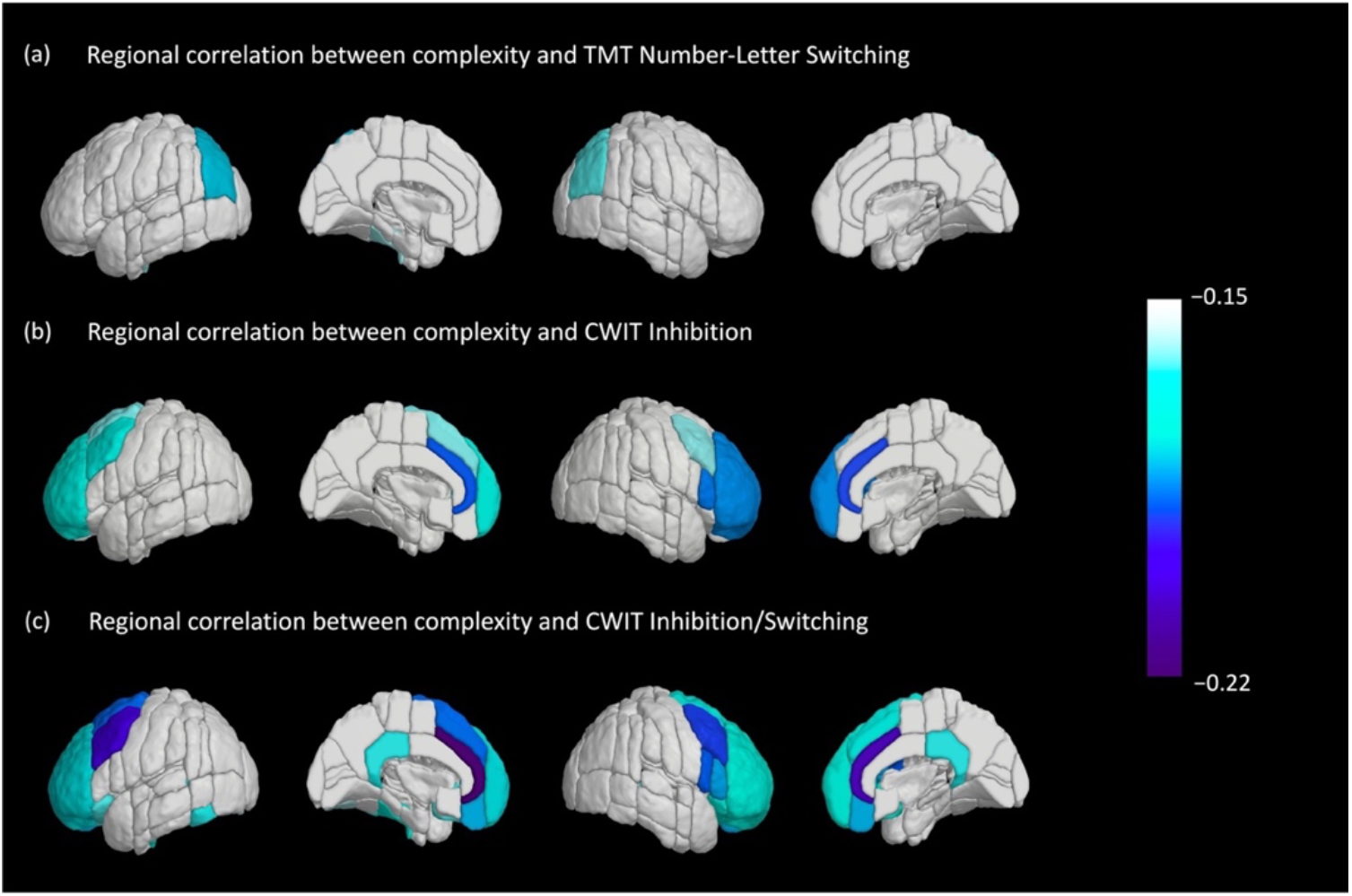
Spearman’s rank correlation coefficients between regional complexity and D-KEFS scores, significant after correction for multiple comparisons (adjusted *p* value 0.00047). (a) Correlation between complexity and TMT Number-Letter Switching. (b) Correlation between complexity and CWIT Inhibition. (c) Correlation between complexity and CWIT Inhibition/Switching.

**Figure 6.**
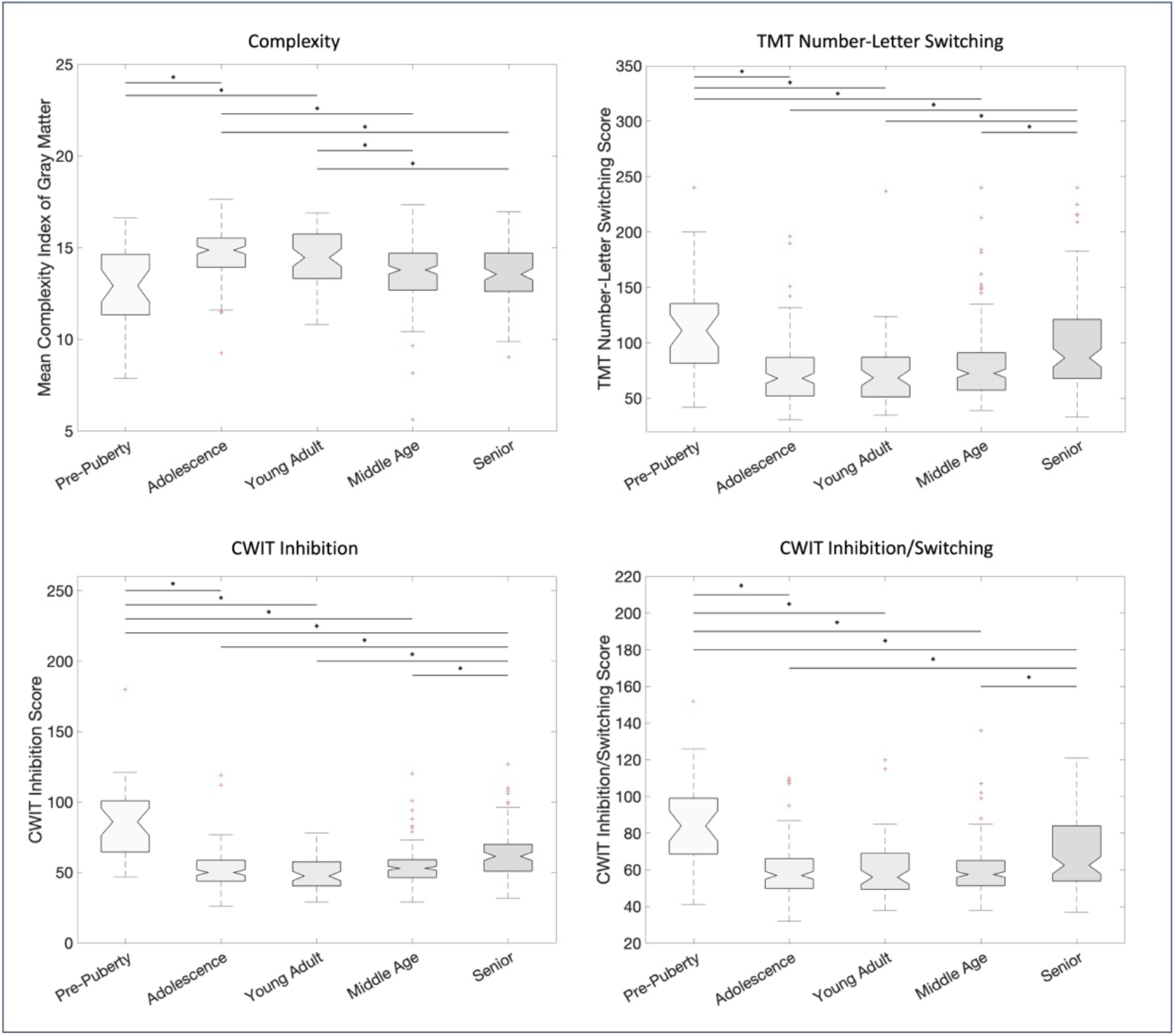
Multiple comparison tests among age groups (a) Complexity vs age (b) TMT Number-Letter Switching score vs age (c) CWIT Inhibition score vs age (d) CWIT Inhibition/Switching score vs age. (* statistically significant age group pairs (*p* value 0.05))

Interestingly, CWIT Inhibition/Switching also showed correlation in posterior cingulate cortex and temporal lobe.

There is an inverted trajectory in D-KEFS test scores across age groups compared to the trajectory of complexity across age groups. The complexity peak maturation in global, female, and male groups occurs in adolescence and young adult age groups whereas lowest time recorded in completing tasks in three D-KEFS tests also associated with adolescence and young adult age group.

## 4 Discussion

In this study we present a comprehensive characterization of cortical lifespan trajectories of MRI-complexity across the human lifespan using MSE. The main observations of this study are that fMRI-complexity shows rapid increase in childhood and adolescence, reaching a peak in early adulthood and then a steady slow decline into late life (Figure 1). This global pattern can be observed in female and male cohorts with a notable maturational delay in male participants reaching peak complexity at a later age than females. A more detailed investigation of complexity in individual cortical areas revealed a general posterior-to-anterior pattern of maturation with frontal areas reaching peak complexity at a later stage (Figure 2). This observation is generally consistent between female and male groups with some marked differences. Another observation is that after reaching the peak, complexity declines more rapidly in parietal and temporal areas, followed by areas in the pre(frontal) cortex. This decline appears to be accelerated in female participants as compared to male participants (Figure 4). However, sex-specific increase and decrease of complexity was not statistically significant. Finally, we also demonstrate that functional complexity is associated with cognitive performance, globally (Table 2), and more specifically executive function in frontal and parietal cortices (Figure 5).

### 4.1 Spatiotemporal cortical trajectories of functional complexity

Our study using fMRI-complexity agrees well with current hypotheses on cortical development, maturation, and aging. Our findings corroborate previous histological and neuroimaging studies that show spatially varying patterns of cortical maturation that may reflect unique developmental processes of cytoarchitectonically determined regional patterns of change. Specifically, the existence of an anatomically defined hierarchical posterior-to-anterior axis of cortical development has been supported by various imaging modalities such as electrophysiology, neuroimaging, and histology ^[34]^. This axis describes the sensorimotor-association cortical axis (S-A axis) of cortical function transition from lower-order sensorimotor to highest-order associative functions. While primary sensory and motor function develop early, the association cortex which supports executive function of the brain mature well into the second decade of life ^[34]^. The dendritic structure of cortical areas is known to undergo significant changes during early maturation including increases in dendritic density and synapse number. Furthermore, during neurodevelopment, pruning of dendritic trees is a critical process associated with specialization and efficiency of brain circuits. Alterations to synaptic and dendritic processes are also central to the pathologies of Alzheimer’s disease (AD). Therefore, the microstructure architecture critically determines the electrical communications of neurons and shape how synaptic signals integrate and support neuronal function. A recent study by Lynch et al observed the spatiotemporal patterns of cortical microstructural maturation in children and adolescents using diffusion MRI. They reported terminal maturation of neurite density index (NDI) in late adolescence for the primary sensory (20.96 years), occipitoparietal (18.21 years), frontotemporal (23.95 years) and limbic (19.36 years) ^[32]^. Besides support from imaging studies for this developmental axis, a study by Gauvrit et al, analyzed the human behavioral complexity over the lifespan using random item generation tasks and observed a peak at age 25 for cognitive function ^[35]^.

Only a few studies have also tried to assess lifespan changes in functional complexity using alternative mathematical descriptions of complexity. One study used Hurst Exponents in a subsample of the NKI cohort (N=116) age ranging from 19 to 85 years and found decrease in complexity with age in frontal and parietal lobes, but they also reported increase in other areas including limbic and occipital brain areas ^[10]^. Another study by Sokunbi et al used Fuzzy Approximate Entropy (fApEn) and SampEn in a cohort of 86 healthy adults with age ranging from 19 to 85 years and reported decline of both complexity metrics with age in frontal and parietal areas ^[9]^. Notably, both studies did not include datapoints from childhood or adolescence and exclusively modeled linear decline of complexity with age. In contrast, our study was able to distinguish maturation as well as decline of cortical function and estimation of the peak age. Another study by Niu et al included participants of a similar age range (N = 319, age range from 8-85 years) as this study and analyzed rs-fMRI complexity using Permutation Entropy (PermEn) ^[11]^. They observed that global PermEn followed an inverted U-shaped trajectory that peaked at approximately age 40 with variation of peak age in individual functional networks ranging between 25 to 52 years. Compared to literature findings of brain maturation, our results are in closer agreement with peak maturation in early adulthood with respect to cortical morphology as well as cognitive function. As a point of notion however, while (Multiscale) SampEn has been most widely used in characterizing neurodevelopmental disorders ^[12,14]^ or alterations in aging and neurodegeneration ^[11,13,15,36,37]^, there are various other mathematical definitions of complexity and the reliability and sensitivity of different complexity methods for fMRI timeseries requires further systematic evaluation. For example, a study comparing different entropy methods observed stable entropy curved for all entropy methods in simulated timeseries data based on 3 distributions except SampEn in Normal and Weibull distributions ^[38]^. However, ApEn, Dispersion Entropy, and PermEn failed to identify differences in brain regions between healthy subjects and subjects with schizophrenia, bipolar and attention-deficit/hyperactivity disorder (ADHD) ^[38]^.

Overall, our study is the first to characterize lifespan trajectories of complexity using MSE and thus expands the repertoire to study cortical development and aging with a metric that assesses functional complexity of the cortex, which complements existing marker of brain structural morphology such as cortical thickness, volume, microstructure, or myelination.

### 4.2 Sex differences in trajectories of functional complexity

Based on the global trajectories we observed that males show a later peak compared to females, which was also replicated in the regional analysis. However, this sex difference is not statistically significant. In the regional analyses separated into male and female participants, the male group showed later peak ages in widespread cortical areas. The cluster analysis further exemplified this observation: in the female group motor temporal cluster was below 6 years whereas in males it was at 25-28 years and the frontal parietal cluster was in the range of 19-24 years in females compared to males it was at 21-24.9 years.

While the cluster analysis showed that the pattern of brain areas with higher or lower rates of increase across the cortex was similar for the whole, female, and male groups, showing highest rates of increase in frontal brain areas whereas occipital and temporal areas showed lower rates, the male group showed generally a higher rate of increase. The sex differences in the rate of increase of complexity as well as in rate of decline are not statistically significant.

### 4.3 Association of functional complexity with cognition

Aiming to find associations between fMRI-complexity and cognitive function we performed correlation analysis for global and regional complexity with D-KEFS test scores. We found that global complexity was inversely correlated with the CWIT and TMT tests. CWIT a neuropsychological assessment tool used to measure cognitive abilities, particularly executive functions like processing speed, inhibition, and cognitive flexibility. A higher CWIT score generally indicates slower processing speed or greater difficulty with interference tasks. TMT requires alternating between numbers and letters, which measures cognitive flexibility and set-shifting abilities. A slower completion time can be indicative of cognitive deficits. For both tests we expected to see inverse relationships with fMRI-complexity since we hypothesized that with higher complexity cognitive functions improve during maturation and with increasing age, fMRI complexity declines along with decline in cognition among healthy subjects. A previous rs-fMRI study by Omidvarnia et al also found that MSE generally correlates with higher-order cognition ^[19]^. Hence our results were in alignment with our hypotheses and expectations, demonstrating fMRI complexity as a marker of functional cognitive abilities. The regional analysis for TMT and CWIT revealed that TMT is primarily associated with parietal lobe and medial temporal lobe whereas CWIT shows effects in lateral and medial frontal cortex and CWIT also includes posterior cingulate cortex and inferior temporal areas. The frontal, parietal and temporal lobes play crucial roles in proper executive function and alterations in these brain areas are linked to behavioral and cognitive deficits in neurodevelopmental and neurodegenerative disorders. Using fMRI complexity could be a novel and promising approach to detect and characterize abnormal trajectories and holds potential as an early marker of subsequent cognitive decline ^[37]^.

### 4.4 Limitations

There are a few limitations to consider when interpreting the findings. First, the data is cross-sectional and to truly assess lifespan trajectories longitudinal data would be ideal. While obtaining data spanning several decades of life is hardly feasible, having a few years of longitudinal data per subjects could improve model trajectories. Second, our data had skewed age distribution in the male cohort. The middle age and senior age groups have lower number of male participants compared to female group. Similar effect can be seen in complexity vs D-KEFS test analysis where there are 314 female participants and 180 male participants. This unequal sample sizes can have an impact on results of the study. Furthermore, there are limitations associated with performing D-KEFS tests such as uncertainties involved with time-dependent measures related to executive function tasks ^[39]^.

### 4.5 Conclusions

Our results indicate that early maturation and slow decline of brain activity can be measured using fMRI complexity. Furthermore, fMRI complexity showed global and regional associations with executive function. Using fMRI complexity could provide a novel and promising approach to detect and characterize abnormal trajectories in atypical neurodevelopment and holds potential as an early marker for cognitive decline in the progression of Alzheimer’s Disease ^[37,40]^.

## Supporting information

see supplemental material

## Acknowledgements

We would like to thank the Nathan Klein Institute for open access neuroimaging data and granting access to phenotypic data of the NKI-Rockland Sample.

Research reported in this publication was supported by the Office of The Director, National Institutes of Health of the National Institutes of Health under Award Numbers R01-AG066711 and S10OD032285. The content is solely the responsibility of the authors and does not necessarily represent the official views of the National Institutes of Health.

## CONSENT STATEMENT

All participants that were enrolled in NKI provided written informed consent.

## DATA AVAILABILITY STATEMENT

Data used in this project is governed by the Nathan Kline Institute and data can be accessed upon request.

