## Supplementary material for "Lifespan Trajectories of the Brain’s Functional Complexity Characterized by Multiscale Sample Entropy": see supplemental material

Age distribution of the sample

Initially 526 subjects with both resting state fMRI and T1 images were chosen randomly from the NKI sample. Twenty-two subjects with composite motion score higher than 0.5 mm were removed from the analysis. The remaining 504 subjects included in the study considered 321 female and 183 male participants. Figure S1 displays the age distribution for the whole study cohorts and female and male groups, respectively.

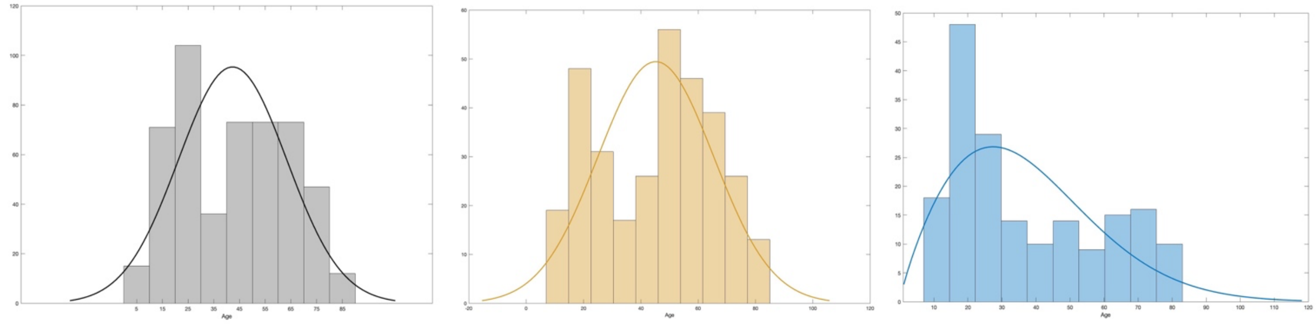

(a) (b) (c)

**S1**: Age distribution of the sample. (a) Age distribution of the whole sample. A Gaussian distribution is fitted to the data ($\mu = 42.3135$, $\sigma = 21.086$). (b) Age distribution of the female group with a fitted Gaussian distribution ($\mu = 42.3135$, $\sigma = 21.086$). (c) Age distribution of the male group with a fitted Weibull distribution ($\alpha = 42.1254$, $\beta = 1.8322$).

Effect of gray matter volume on complexity analysis

Gray matter volume of each subject was calculated using FMRIB Software Library (FSL). By applying FSL command "fast" on T1 images, images were segmented into tissue compartments and "fslstats" command was used for tissue volume quantification. This provides partial volume estimates for the volume quantification. Gray matter volume decreases with age. This can influence the results obtained for the mean gray matter complexity vs age distribution. Therefore, to evaluate the influence of gray matter volume in our analysis, a generalized linear regression model (MATLAB 'fitglm') was applied among complexity, age and mean gray matter volume as a covariate (Table S1).

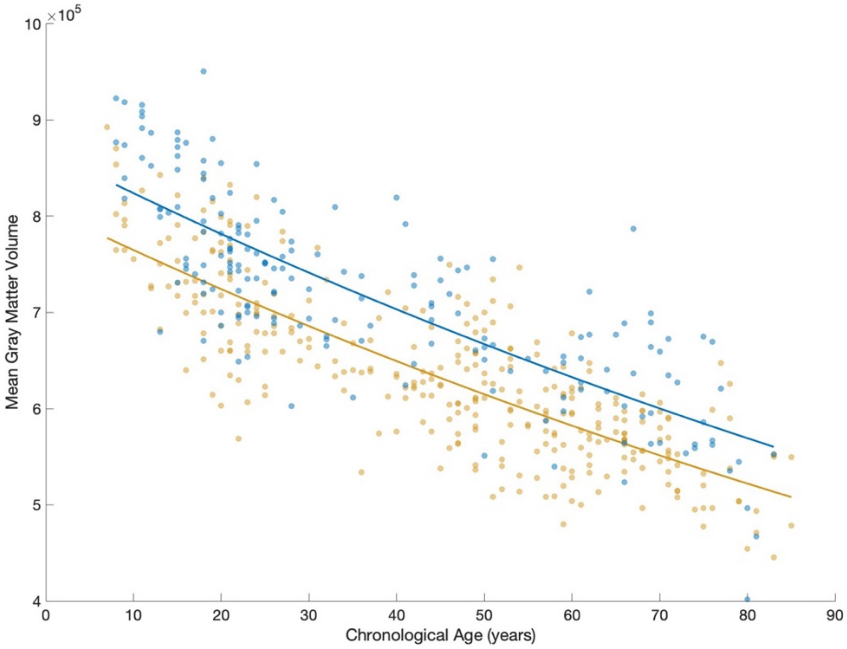


**S2**: Gray matter volume vs chronological age in years (female in yellow and male in blue).

$$Mean\ gray\ matter\ complexity \sim 1 + \ Age + Mean\_graymatter\_volume$$

**Table S1**: Results of the generalized linear regression model

| | Estimate | SE | t-statistic | P value |
|---|---|---|---|---|
| Age | $-0.015478$ | $0.005586$ | $-2.7708$ | $0.0058$ |
| Mean Gray matter Volume | $-6.0477 \times 10^{-9}$ | $1.1824 \times 10^{-6}$ | $-0.0051$ | $0.9959$ |

*Standard Error (SE)

Bayesian Information Criterion (BIC) for model fitting

For the ROI analysis of 105 regions, we applied five functions for each ROI to evaluate the lowest BIC. For the total cohort we observed 6 regions with lowest BIC values for linear fit, 31 regions with quartic fit and 68 regions with double exponential fit. We observed no region with quadratic and cubic polynomial fits. For female subjects we observed 46 regions with linear fit, 3 regions with quadratic fit, 9 regions with quartic fit and 47 regions with double exponential fit. For male subjects there were 7 regions with linear fit, two regions with cubic and quartic fits and 96 regions with double exponential fit (Table S2). Among three categories of the analysis (global, female, and male), most ROIs exhibited a double exponential trajectory depicting an initial increase in complexity followed by a decrease after a peak. Contrary, regions that have linear association with age show primarily a decrease in complexity after the age of 6 years.

**Table S2**: Number of ROIs for each polynomial fit

| **Polynomial** | Number of ROIs (C index - Global) | Number of ROIs (C index - Female) | Number of ROIs (C index - Male) |
|---|---|---|---|
| Linear | 6 | 46 | 7 |
| Quadratic | 0 | 3 | 0 |
| Cubic | 0 | 0 | 1 |
| Quartic | 31 | 9 | 1 |
| Double Exponential (Exp (2)) | 68 | 47 | 96 |

In figure S3 we display (a) exemplary trajectories for different models and (b), (c), and (d) the best model for each cortical area for each of the grouped analyses.

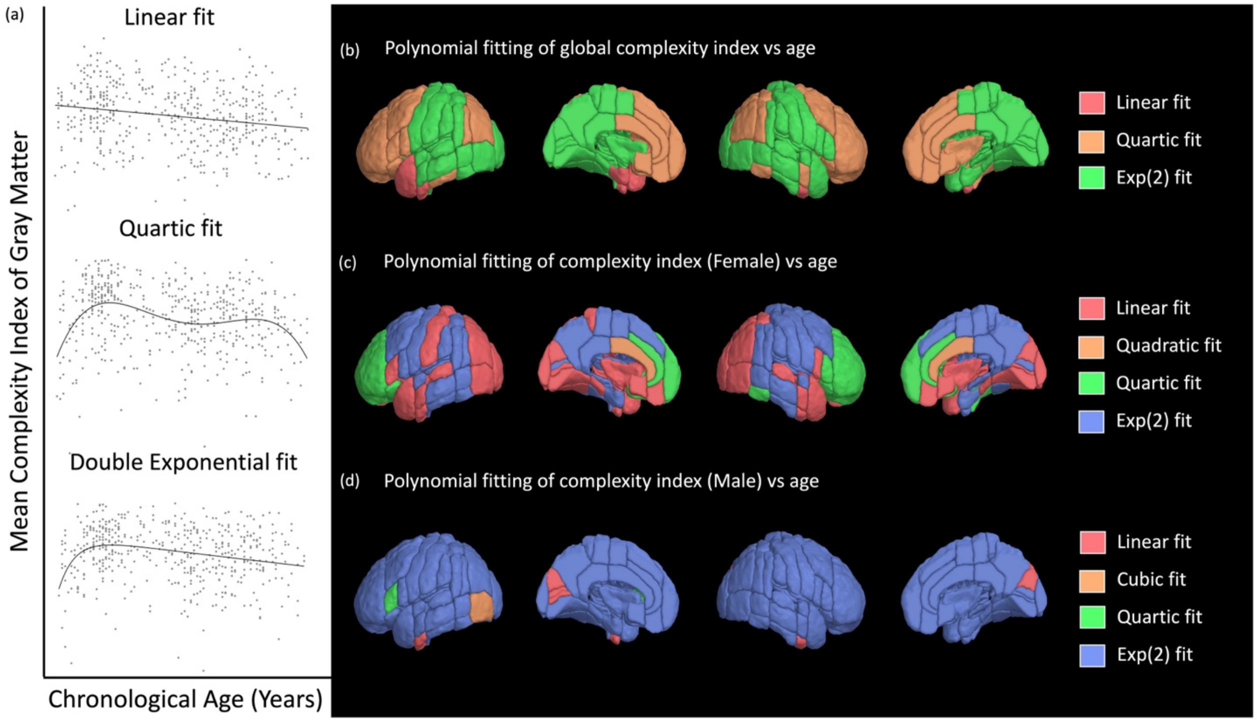


**S3**: Polynomial fitting with Bayesian Information Criterion (BIC). (a) Linear, quartic, and double exponential fit for temporal pole (left), frontal pole (right), and superior frontal gyrus (right) for global complexity index. (b) Polynomial fitting in global complexity index vs age. (c) Polynomial fitting in complexity index (female) vs age. (d) Polynomial fitting in complexity index (male) vs age.

Delis-Kaplan Executive Function System test scores (D-KEFS) along the lifespan

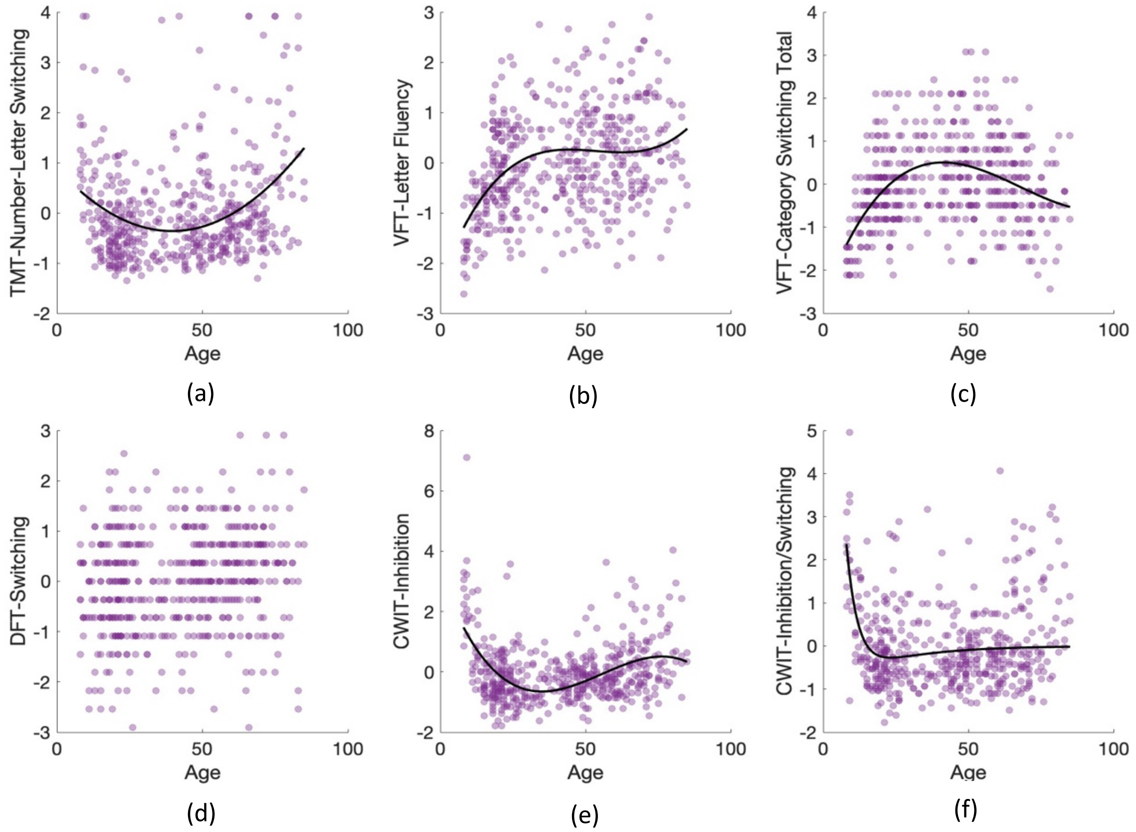


**S4**: D-KEFS test scores vs Age. Bayesian Information Criterion (BIC) was used to choose the best polynomial. (a) TMT Number-Letter Switching vs age fitted with a quadratic polynomial. (b) VFT Letter fluency vs age, (c) VFT Category Switching scores vs age, (e) CWIT Inhibition vs age and (f) CWIT Inhibition/Switching vs age fitted with cubic polynomials. (d) DFT Switching vs age. Polynomial fitting is not applied on DFT Switching vs age distribution.

Mean Gray Matter Complexity vs Delis-Kaplan Executive Function System test scores (D-KEFS)

Correlation coefficients between mean gray matter complexity and D-KEFS tests were calculated using both Spearman's rank correlation coefficient and Pearson correlation coefficient.

| D-KEFS Test | Pearson Correlation | P value |
|---|---|---|
| TMT<br>Number-Letter Switching | -0.1262 | 0.0050* |
| VFT<br>Letter Fluency | -0.0011 | 0.9811 |
| VFT<br>Category Switching Total | 0.0136 | 0.7629 |
| CWIT<br>Inhibition | -0.1200 | 0.0076* |
| CWIT<br>Inhibition/Switching | -0.1787 | 0.0000648* |
| DFT<br>Switching | 0.0301 | 0.5051 |

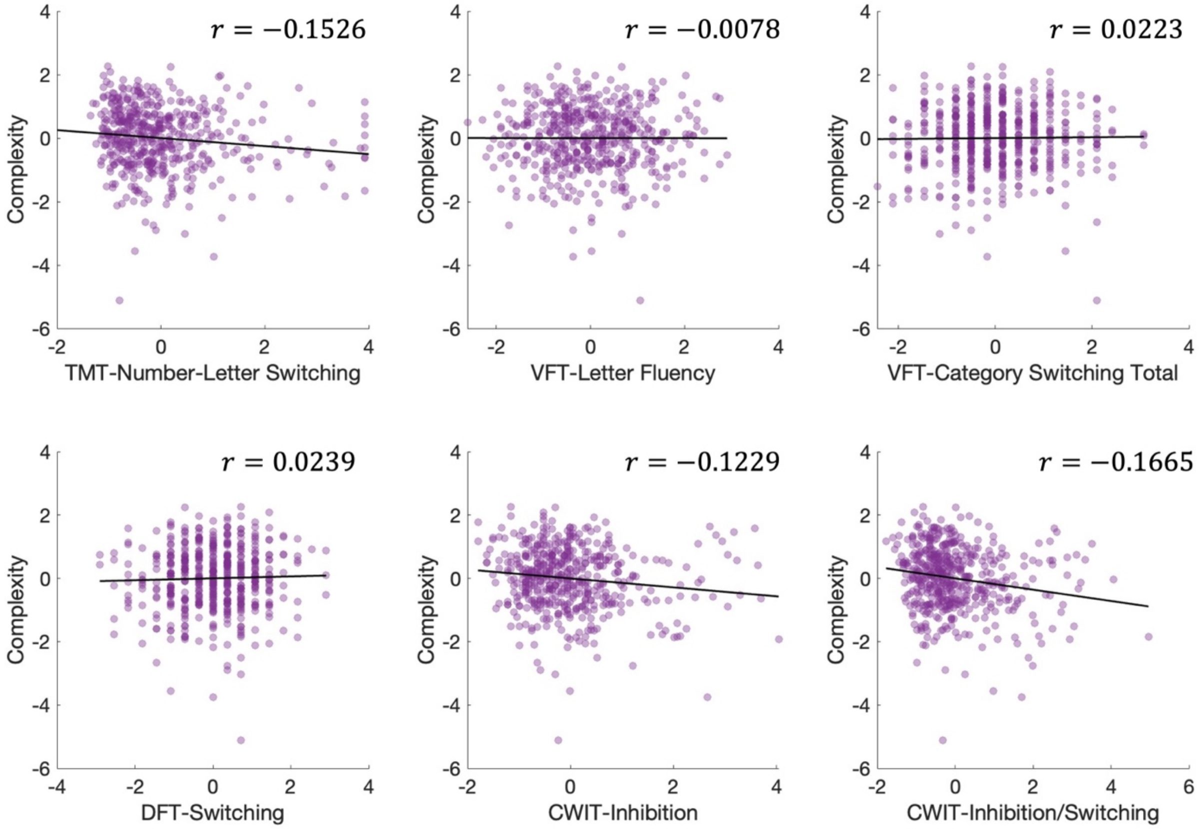


**S5**: Mean gray matter complexity vs D-KEFS test scores. Spearman's rank correlation coefficients are displayed on each figure panel.
